# Photographic age gap is associated with frailty and predicts survival in laboratory mice

**DOI:** 10.64898/2026.11.02.762463

**Authors:** Kevin Perez, Clement Molina, Luca Tagliafico, Feng Ruiqi, Kamil Pabis, Miles Sarill, Giorgia Bigossi, Serena Marcozzi, Amirhassan Monajemi, Andrea Alimonti, Fabrizio d’Adda di Fagagna, Dmitry Bulavin, Marco Malavolta

## Abstract

Ageing studies in mice need biomarkers that are cheap, non-invasive and repeatable. We asked how much of a mouse’s age can be read from a simple photograph, and whether the part that cannot be explained by age carries biological information. We assembled 2,846 photographs of 1,292 mice (four strains, 1 to 34 months old) from seven facilities and predicted age with a frozen self-supervised vision model (DINOv2) and a ridge regression, always tested on facilities the model had never seen. Absolute age transferred poorly between facilities (median absolute error 138 days), because each facility’s photographs carried an offset. Relative age transferred well: among mice photographed in the same session, the older mouse was predicted older in 65% of pairs, rising above 95% for age differences over 500 days, and the later photograph of the same mouse was predicted older in 63% of repeat pairs. Sex, in comparison, was inferred more reliably (87%). Mice predicted older than their age were frailer (+0.12 SD per 100 days) and had shorter survival (hazard ratio 1.14 per 100 days), and fourth generation (G4) telomerase-deficient mice looked 151 days older than their wild-type controls. Simple photographs therefore carry a real biological age signal that can be read quickly and cheaply, best used to compare mice photographed together.

## Introduction

Testing interventions on ageing in mice needs a read-out of biological age that can be taken often, cheaply and without harm. Lifespan is the definitive end point but arrives late, and molecular clocks need tissue or blood and specialised assays [1–3]. Clinical frailty indices are non-invasive and predict survival in mice [4–6], but they take trained staff and vary between scorers; machine vision can now score frailty from video of freely moving mice [7]. Photographs are often taken to illustrate the effect of potential anti-ageing treatment, but these are never reliably quantified [8–10].

In humans, facial photographs predict chronological age and carry information on health and survival beyond it [11–13]. Whether the same holds for mice is unknown. Mice are small, often photographed in different poses and settings, and their visible ageing signs — coat greying and thinning, posture, body shape, eye changes — are subtler than in humans [4].

Here we collected photographs from seven facilities and asked several questions. How much of a mouse’s age can be read from a photograph of a facility the model has never seen [14]? Can the model rank younger and older mice? Does the same mouse photographed over time look older as it ages? How does the performance compare to a human expert prediction? Lastly, does the difference between photographed and real age — the photographic age gap — relate to frailty, survival, sex, strain and a genetic model of accelerated ageing? We also use sex prediction as a comparator for what signal photographs can carry.

## Results

### A multi-facility photograph collection and a frozen-feature age model

The dataset holds 2,846 photographs of 1,292 mice from seven facilities (Fig. 1). Facilities differed in number of mice (19 to 692), age range covered, strain, sex balance, camera, background and pose (Fig. 1b–e). Four strains were represented: C57BL/6J (963 mice), BALB/c (182), genetically heterogeneous UM-HET3 [15] (130) and telomerase-deficient *Terc*⁻/⁻ mice of the third (G3) and fourth generation (G4) [16] (TERC⁻/⁻ G3, 11; TERC⁻/⁻ G4, 6). Two facilities photographed only males. 235 mice were photographed at two or more ages.

**Fig. 1.**
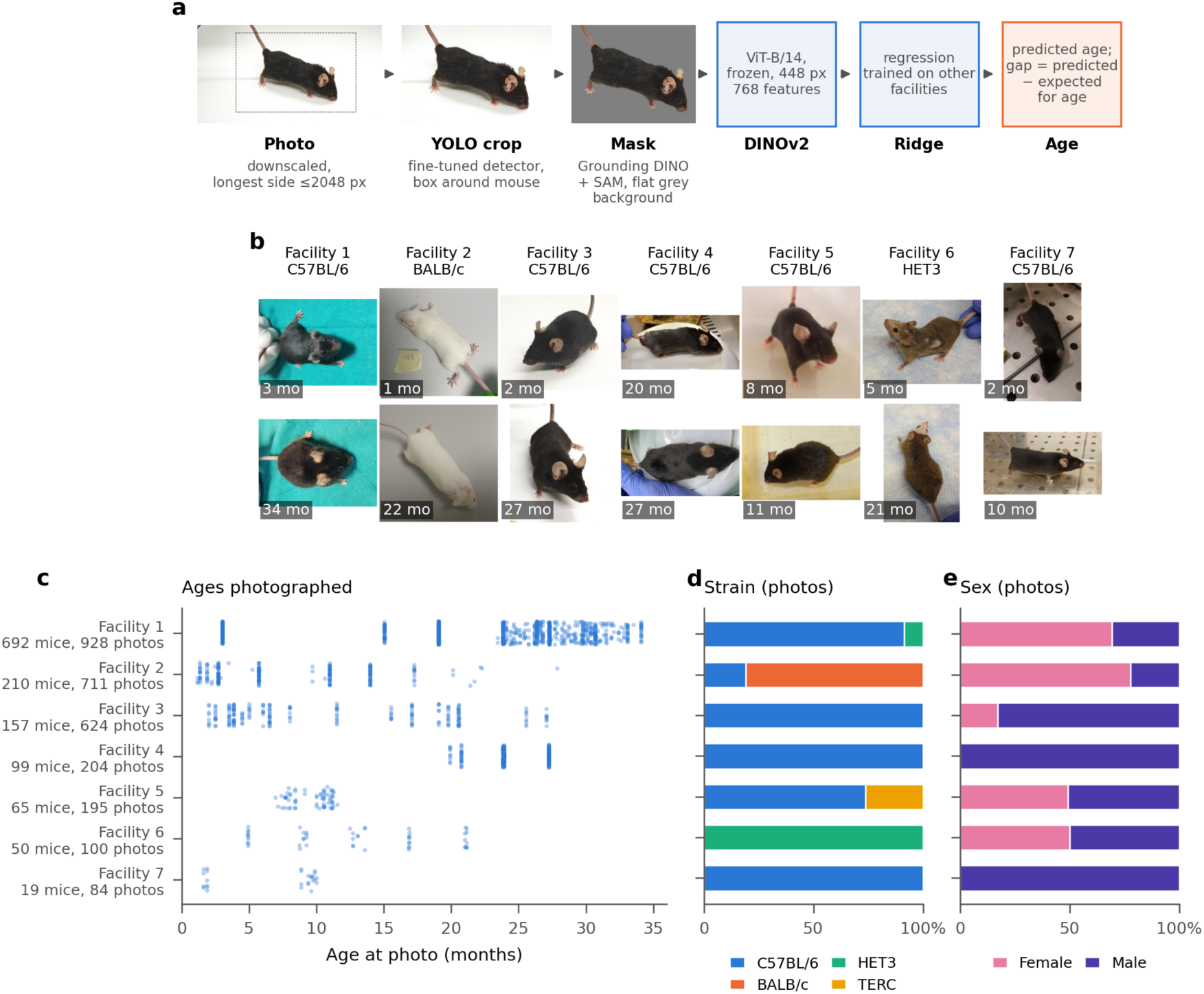
Dataset and pipeline. **a**, Processing of one photograph: downscaling, YOLO crop, background removal with Grounding DINO and SAM, frozen DINOv2 embedding and ridge regression; the photographic age gap is the predicted minus the expected prediction for a mouse of that age. **b**, Example photographs from each facility before background removal (YOLO crops, shown whole), youngest (top) and oldest (bottom) control mouse of the facility’s main strain. **c**, Ages photographed per facility (one dot per mouse and session), with numbers of mice and photographs. **d**,**e**, Share of photographs by strain and by sex.

Each photograph was downscaled, the mouse was located by a fine-tuned YOLO detector [17] and cropped around its detected box, and its outline was traced by Grounding DINO [18] and SAM [19] so that the background could be replaced by flat grey (Fig. 1a). A self-supervised vision transformer (DINOv2 ViT-B/14 [20]), used without any training on mice, turned each crop into 768 numbers, and a ridge regression mapped these to age. We chose this frozen-feature design over fine-tuning the network, because fine-tuning did not transfer to new facilities (see below). Unless stated otherwise, every prediction was made by a model fitted without the photographed mouse’s facility (leave-one-facility-out), and photographs of the same mouse in the same session were averaged (1,530 mouse-sessions of control mice).

### Relative age transfers to new facilities; absolute age does not

In facilities the model had never seen, predicted and true age correlated (r = 0.74; Fig. 2a), but the mean absolute error was 162 days (median 138 days). That is better than guessing the overall median age (243 days) or the mean age of each photographing session (361 days), but too large for an absolute clock. Most of the error came from facility-level offsets: one facility’s mice were predicted 276 days too young on average and another’s 176 days too old (Fig. 2f). When the model had seen other mice of the same facility (leave-one-mouse-out), the error fell to 93 days (r = 0.91).

**Fig. 2.**
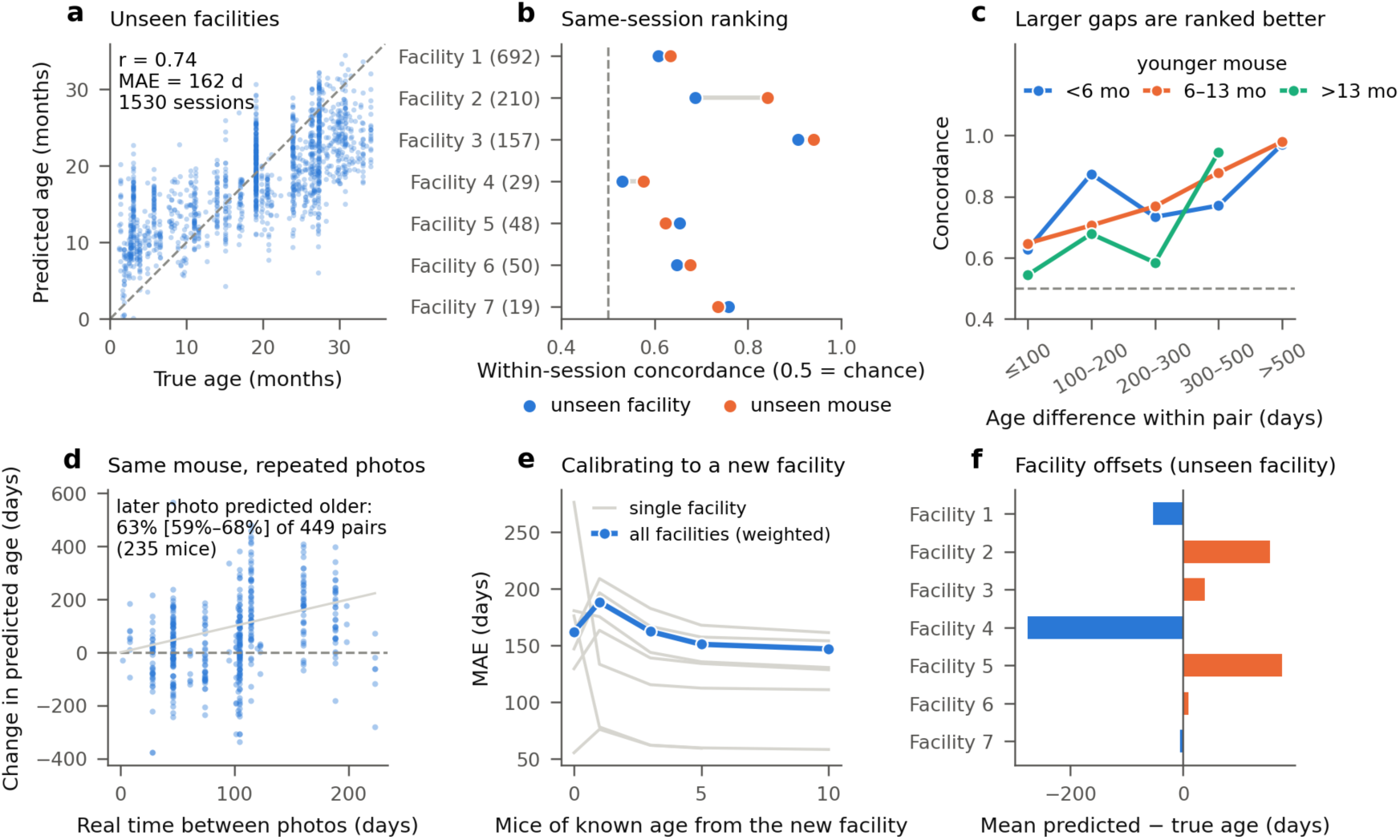
Age prediction in facilities the model has never seen. **a**, Predicted against true age (leave-one-facility-out; one dot per mouse and session). **b**, Within-session concordance per facility (numbers of mice in brackets), for unseen facilities and unseen mice; dashed line, chance. **c**, Concordance by age difference within a pair, by age of the younger mouse (bins with ≥50 pairs). **d**, Same mouse photographed twice: change in predicted against real time; grey line, perfect agreement. **e**, Error after re-centring a new facility’s predictions on k mice of known age (grey, single facilities; blue, all facilities weighted). **f**, Mean offset (predicted minus true age) per unseen facility.

We therefore measured what does not depend on the offset: the ranking of mice photographed in the same session. For every pair of mice of different age photographed together, we asked whether the older one was predicted older (0.5 = chance). Within-session concordance was 0.65 (95% CI 0.62–0.68) for unseen facilities and 0.69 (0.66–0.72) for unseen mice (Fig. 2b). It ranged from 0.53 in the facility whose mice were all 20 to 27 months old to 0.91 in a facility spanning 2 to 27 months. Concordance depended on how far apart the two mice were in age: about 0.54–0.65 for differences under 100 days and 0.97–0.98 above 500 days, whatever the age of the younger mouse (Fig. 2c).

The same held within individual mice. Of 449 pairs of photographs of the same mouse at two ages (235 mice, 91% of pairs taken in different sessions), the later photograph was predicted older in 63% (59–68%; Fig. 2d). Predicted change was smaller than real change (median 0.53 predicted days per real day).

Knowing the age of a few mice from a new facility reduced its offset. Re-centring predictions on 5 or 10 mice of known age lowered the error from 162 to 151 and 147 days; a single mouse made it worse (188 days), because one mouse’s own error is as large as the offset (Fig. 2e).

Two design choices were tested and are reported for transparency. Using 448-pixel instead of 224-pixel input lowered the error by 7.6 days (4.0–11.1) without changing concordance (+0.009, −0.011 to 0.028; Supplementary Fig. 1); we kept 448 pixels for that reason alone. Fine-tuning the network itself on age, the usual way to adapt such a model, did not transfer to new facilities. We tried training the whole network end to end, and a more conservative version that started from the ridge solution and kept the early layers frozen; neither matched the frozen model on held-out facilities, and end-to-end training collapsed to near-constant predictions in the largest facility. We therefore report the frozen model throughout.

### Sex is predicted more easily than age, and the model ranks age where human experts cannot

To know what to expect from a photograph, we trained the same model to predict sex, with the same leave-one-facility-out design. Using the same within-session measure, the male of a male–female pair was ranked as more male in 87% of pairs (84–90%), against 66% (63–69%) for age in the same run (Fig. 3a). In two facilities sex was read almost perfectly (96–97%). Sex is fixed and visible in body size and shape; age changes slowly and is confounded with body size only part of the way. The gap between the two shows that the model is not limited by image quality, and that age is the harder signal.

**Fig. 3.**
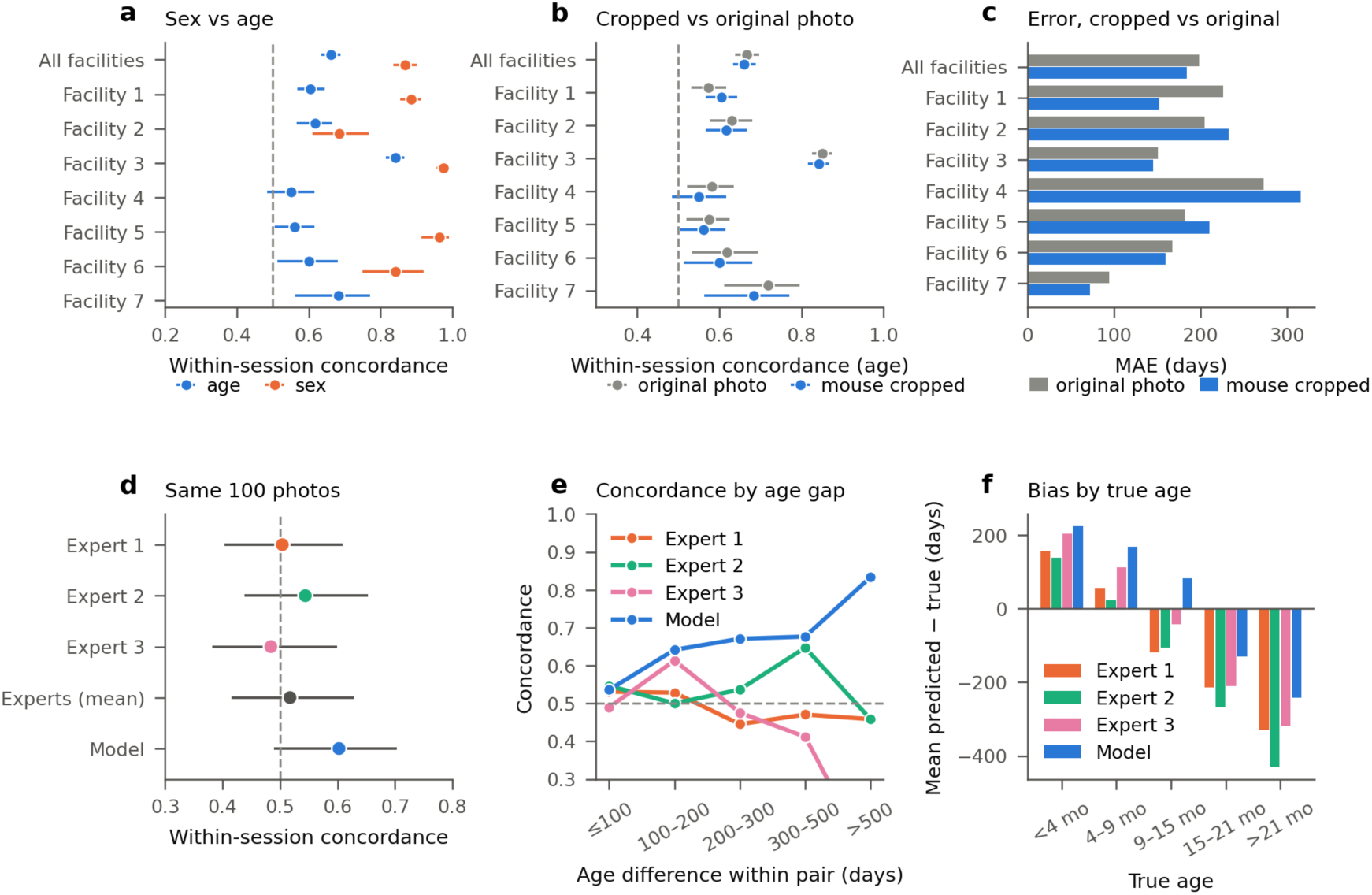
What photographs can and cannot tell. **a**, Within-session concordance for sex (male–female pairs) and age (pairs of different age) from the same model design; bars, 95% CI from resampling mice. **b**,**c**, Concordance and error with the background removed or kept. **d**, Concordance of three expert raters, their averaged estimate and the model on the same 100 photographs. **e**, Concordance by age difference. **f**, Mean bias by true age band.

Removing the background did not change how well mice were ranked. Concordance was 0.66 with the mouse cut out and 0.67 on the YOLO crop with its background kept; the mean error was lower with cropping (184 versus 199 days; Fig. 3b,c). We kept the cropped version because it removes cues such as cage, bedding and hands that differ between facilities without costing accuracy.

Three experienced mouse researchers independently estimated the age of the same 100 photographs from facilities the model had not seen, 10 of them shown twice without notice. Two were self-consistent (Spearman ρ = 0.86 and 0.77 between repeats) and the third less so (0.50); they agreed moderately with each other (ρ = 0.52–0.67; mean difference 3.4–4.4 months), and their mean errors matched the model’s on the same photographs (201, 209 and 212 versus 198 days). Their within-session concordance, however, was at chance (0.50, 0.40–0.61; 0.54, 0.44–0.65; and 0.48, 0.38–0.60; 0.52 for their averaged estimate) and did not rise as age differences grew, whereas the model’s rose from 0.54 for differences under 100 days to 0.83 above 500 days (Fig. 3d,e). All three experts overestimated mice younger than 4 months (by 157, 138 and 204 days) and underestimated mice older than 21 months (by 329, 430 and 319 days), pulling every estimate towards the middle; the model showed the same pattern, stronger in young mice (+224 days) and weaker in old ones (−243 days) (Fig. 3f). On these 100 photographs the model ranked mice better than each of the three raters (concordance 0.60 versus 0.48–0.54), but the difference was not significant (raters’ average minus model −0.09, 95% CI −0.23 to 0.08).

### Ranking works across sexes and strains, but photographic age differs between them

The model ranked mice of each sex and strain above chance (Fig. 4a,c; within-session concordance among pairs of the same group). Concordance was 0.61 (0.58–0.65) among females and 0.75 (0.69–0.80) among males, but this difference mainly came from one facility that photographed mostly males (0.91 among males); within the five facilities with both sexes, neither sex was ranked consistently better. Concordance was similar for C57BL/6J (0.65, 0.61–0.69), BALB/c (0.68, 0.60–0.74) and UM-HET3 mice (0.63, 0.54–0.71).

**Fig. 4.**
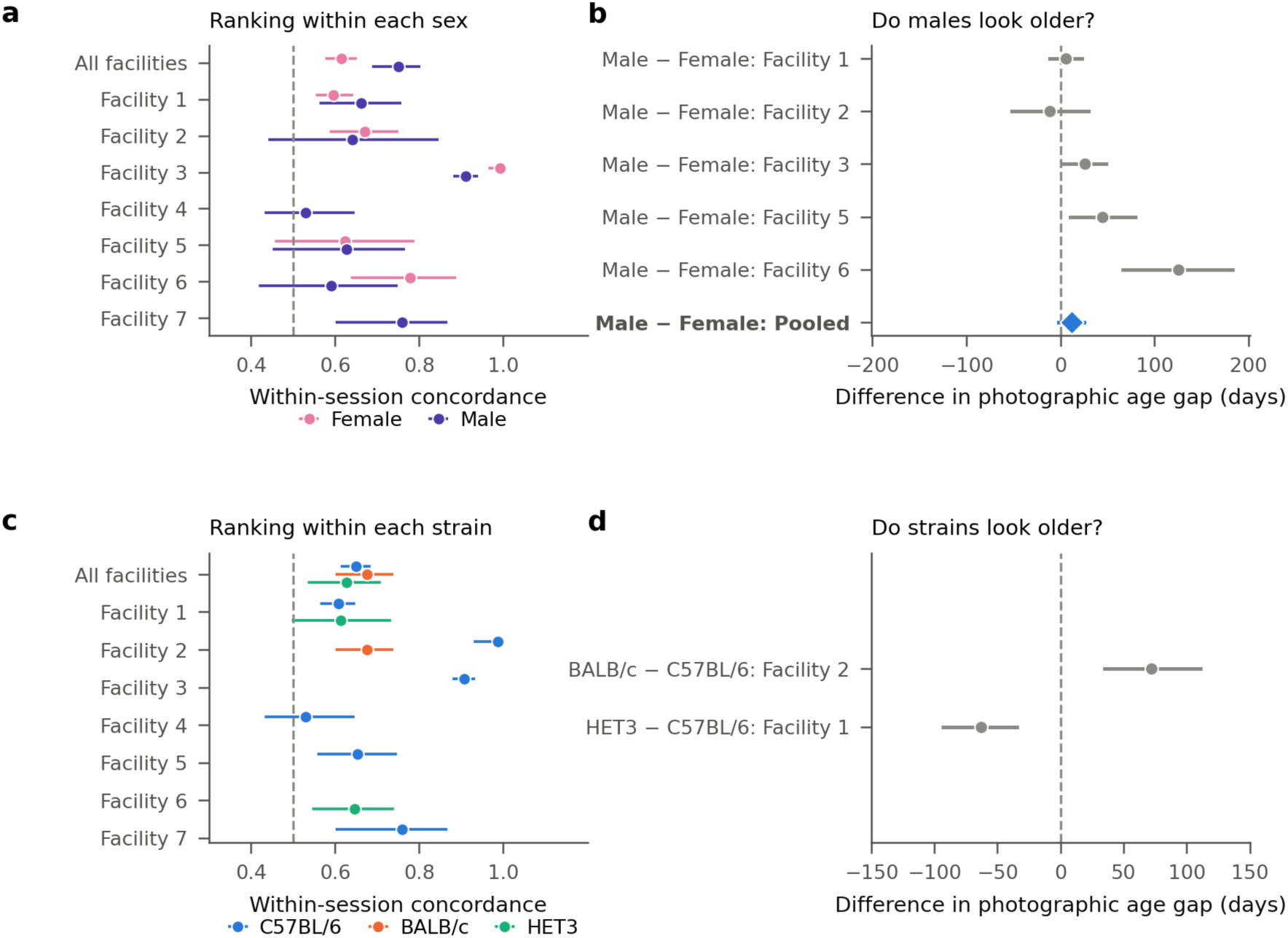
Sex and strain. **a**,**c**, Within-session concordance computed separately within each sex (a) and strain (c), per facility and overall. **b**,**d**, Difference in photographic age gap between males and females (b) and between strains (d) within facilities that contain both groups; diamond, pooled estimate weighted by mice. Bars, 95% bootstrap CI.

The photographic age gap — predicted minus expected age for a mouse of that age in that facility — did differ between groups. Males looked 12 days older than females of the same age, a difference that was not significant (−2 to 25; Fig. 4b); it was positive in four of the five facilities with both sexes (from +6 to +125 days; the fifth −11, −52 to 30). BALB/c mice looked 72 days older than C57BL/6J mice in the same facility (35–111), and UM-HET3 mice 63 days younger (34–93; Fig. 4d).

### Mice that look older are frailer and die sooner

If photographs capture biological and not only chronological age, mice predicted older than their age should be frailer. Two facilities scored frailty at the time of the photograph, one with a vitality score [21] (296 mice) and one with a frailty index [4] (98 mice). We reversed the sign of the vitality score so that higher means frailer in both, adjusted for age within each facility, and expressed the result per 100 days of photographic age gap. Frailty increased by 0.12 SD per 100 days (0.06–0.18; P < 0.001), with almost the same effect in both facilities (0.12 and 0.11 SD; heterogeneity I² = 0; Fig. 5a,e,f). Body weight, available with enough mice in one facility, was not related to the gap (0.04 SD, −0.04 to 0.12).

**Fig. 5.**
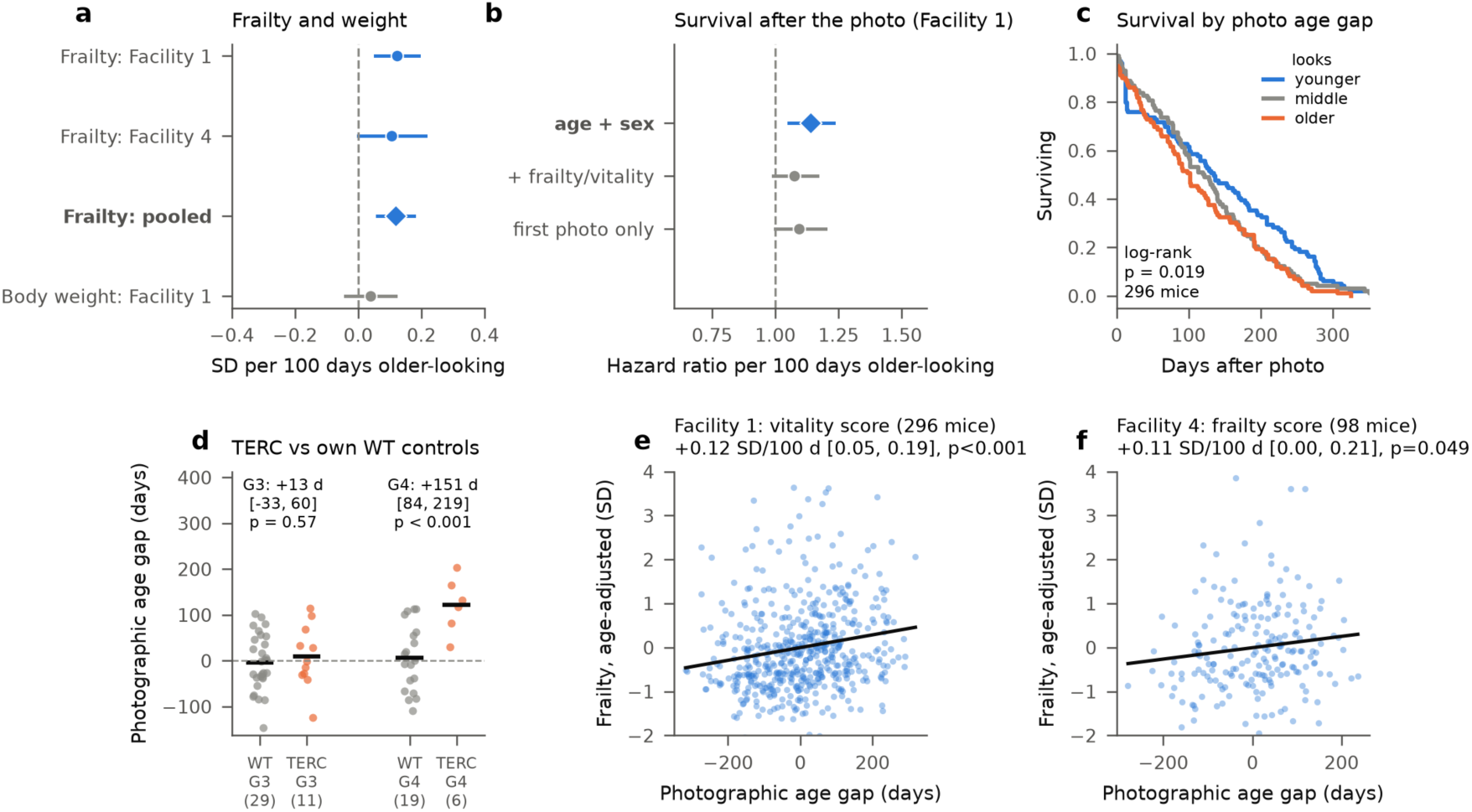
The photographic age gap as a biomarker. **a**, Change in frailty (frailty index, or vitality score with sign reversed; standardised within facility, higher = frailer) and body weight per 100 days of age gap, adjusted for age; diamond, fixed-effect pooled estimate. **b**, Hazard ratio of death per 100 days of age gap, Cox models (Facility 1, control mice). **c**, Survival after the photograph by thirds of age gap (Facility 1, control mice, first photograph; unadjusted illustration). **d**, Age gap of third- and fourth-generation telomerase-deficient mice (TERC⁻/⁻ G3, G4) and the wild-type cohort photographed alongside each; black line, mean; differences adjusted for age. **e**,**f**, Age-adjusted frailty against age gap in each facility; line, least-squares fit; estimate as in a.

Mice were followed until death in both facilities; we analysed survival in the facility with the larger cohort (296 control mice), because the other had only 29 control mice. Adjusting for age and sex, each 100 days of photographic age gap raised the hazard of death by 14% (hazard ratio 1.14, 1.06–1.23; P = 0.001; 296 mice followed to death, 531 photograph sessions; Fig. 5b). The association weakened when the vitality score was added to the model (1.08, 0.99–1.17), which itself predicted death strongly (1.51 per SD), and when only each mouse’s first photograph was used (1.09, 1.00–1.20). Survival differed between thirds of the age gap (log-rank P = 0.019; first photograph of each mouse; Fig. 5c).

Finally, we asked whether a genetic model of accelerated ageing looks older. Each generation of telomerase-deficient mice was compared with the wild-type cohort photographed alongside it. Fourth-generation mice (TERC⁻/⁻ G4), known to have a more severe phenotype and shorter survival [22], looked 151 days older than their wild-type controls (84–219 days; P < 0.001; 6 versus 19 mice), whereas third-generation mice (TERC⁻/⁻ G3) did not differ (+13 days, −33 to 60; P = 0.57; 11 versus 29 mice; Fig. 5d).

### The model reads the whole body, not one feature

To see which parts of a photograph drive the prediction, we covered small patches of each image in turn with flat grey and recorded how much the predicted age changed (occlusion maps [23]; Fig. 6a). In old mice the face and ears often pushed the prediction up, while sharp body fur pushed it down.

**Fig. 6.**
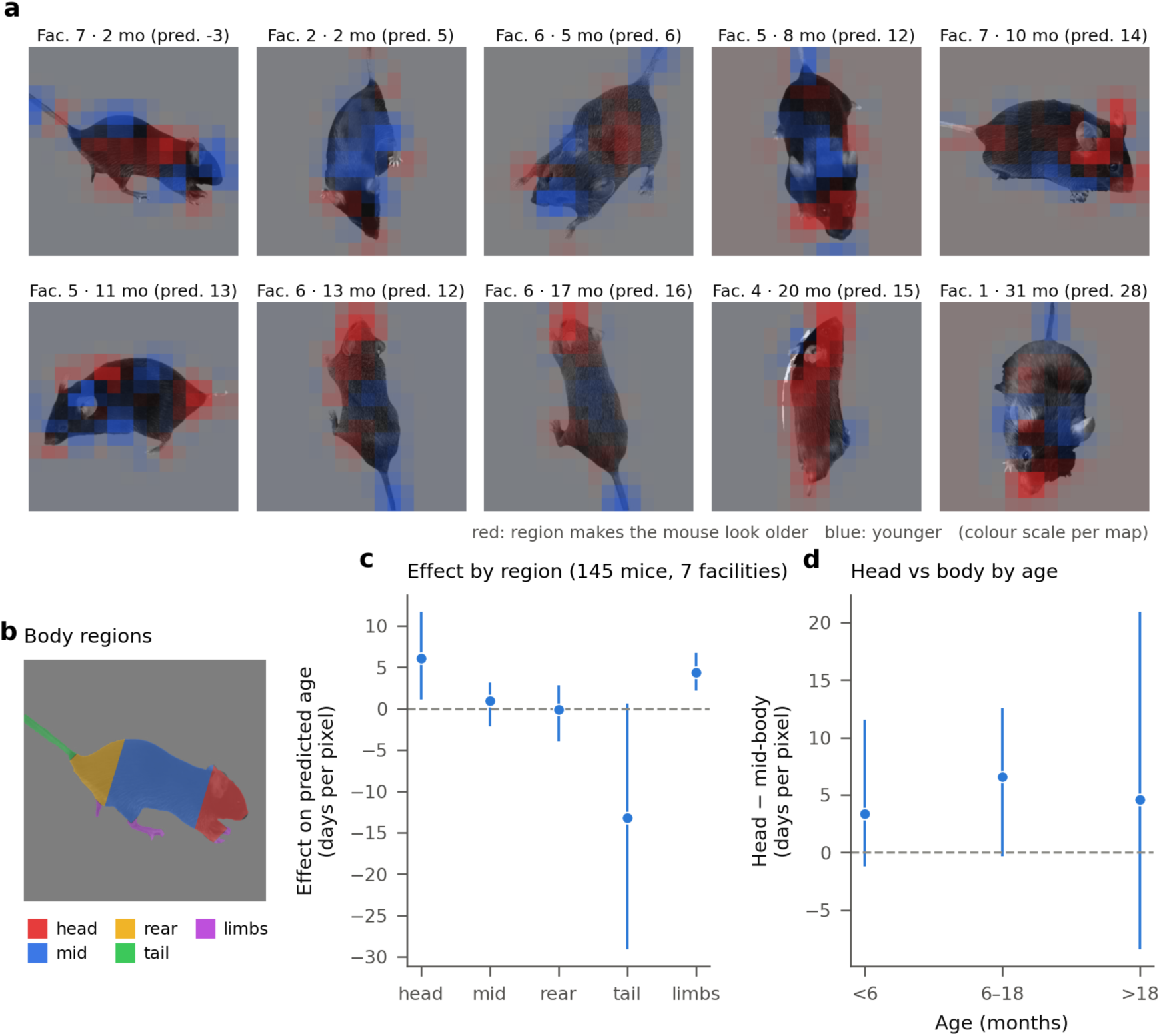
What the model looks at. **a**, Occlusion maps of ten mice from young to old (patches replaced by flat grey; drawn over a greyscale copy of each photograph); red, region raises predicted age; blue, lowers it (scale per map). **b**, Body regions defined from the mouse outline. **c**, Mean effect per pixel of each region (grey occlusion; 145 oriented photographs of 133 mice, 7 facilities). **d**, Head minus mid-body by age group. Bars, 95% CI from resampling facilities.

We quantified this on 221 photographs of 199 mice (about 30 per facility), dividing each mouse into head, mid-body, rear, tail and limbs; regions could be assigned in the 145 photographs (133 mice) in which the tail was visible (Fig. 6b,c). These photographs showed younger mice than the rest (357 versus 504 days, P < 0.001), so region estimates under-represent old mice. The tail region pushed predictions younger (−13 days per pixel; 5 of 7 facilities), and the head (+6) and limbs (+4) older. The head pushed predictions older relative to the mid-body at all ages (Fig. 6d; +3, +7 and +5 days per pixel below 6 months, at 6–18 months and above 18 months), with no trend with age (P = 0.37). No single region dominated: the model draws on the whole body, with the head contributing most consistently.

## Discussion

A single photograph carries a real signal of a mouse’s age, and part of that signal tracks frailty and survival. The signal is relative, not absolute. A model trained elsewhere ranks mice photographed together and follows the same mouse over time, but it cannot give an age estimate in days in a new facility, because each facility’s photographs shift all predictions by up to nine months. This distinction decides how photographic age can be used.

### What photographs can and cannot tell

Three comparisons place the age signal. Sex, which is fixed and visible, was predicted more reliably by the same model; age is a harder target, not one limited by image quality. Three experienced researchers did no better than chance at ranking mice photographed together, so the model captures age cues that even experienced researchers do not use. And removing the background changed nothing, so the model is not reading cages or bedding. The occlusion maps show that the prediction draws on the whole body rather than one feature, as expected if it reads diffuse age-related changes such as greying and thinning fur [4].

### A photographic age gap

Mice that looked older than their age were frailer and died sooner, and fourth-generation telomerase-deficient mice looked older than their wild-type controls. Each 100 days of gap corresponded to 0.12 SD more frailty and a 14% higher hazard of death. The survival association weakened once the vitality score was known, so a photograph captures part of what a frailty assessment measures, from an image that a technician can easily take. Its advantages are cost and repeatability: it needs no trained scorer and can be taken at every handling.

### Pitfalls for image-based clocks

Our results show how important it is to assess clocks in multiple facilities. Tested on mice from facilities it had already seen, the same model cut its error by 43% (93 versus 162 days). Fine-tuning the network made transfer worse and collapsed in the largest facility. Sex and strain shifted photographic age by weeks to months, plausibly through body size and coat colour. Any comparison of groups therefore needs controls photographed in the same session, and claims about absolute age need validation in a facility the model has not seen.

### Limitations

Facilities differed in strain, sex, age range and photographing protocol at the same time, so these effects cannot be fully separated. Frailty scores were available in two facilities, and survival could be analysed in one. The fourth-generation telomerase-deficient group was small (six mice, one facility). Only three human raters were tested, on 100 photographs. Photographs were taken without a standard pose or lighting, so performance under a standard protocol is likely to be higher.

### Outlook

A standard protocol — same pose, background and lighting, with a few age references per session — should reduce the facility offsets that dominate the error. Photographing the same mice at each routine handling would turn the within-mouse signal into a longitudinal read-out at little extra cost. We release the code and models so that facilities can test photographic age on their own mice. We also release the training dataset in anticipation of further improvements in future releases.

## Methods

### Animals and photographs

Photographs were taken for this study at seven facilities under each institution’s approved animal protocols. For each photograph we recorded the mouse identifier, age in days, sex, strain, facility and photographing session (one folder per session). All mice used in training were untreated control mice.

Telomerase-deficient (Terc⁻/⁻) mice were bred at IFOM (Facility 5) from Terc⁺/⁻ mice (B6.Cg-Terc^tm1Rdp/J^, Jackson Laboratory, stock 004132) by successive intercrosses of Terc⁻/⁻ mice up to the third (G3) and fourth (G4) generation [24, 25]. Some G3 and G4 mice had received a control antisense oligonucleotide against an unrelated sequence, whereas others were untreated. These groups were pooled for analysis, as previously reported, because no consistent differences were observed in multiple ageing-related parameters [25]. Telomerase-deficient mice were excluded from training and used only as a test group.

At EPITERNA (Facility 4), some mice had received various treatments; these mice were excluded from training and from the main evaluation and only included in the frailty analysis with treatment effect ignored.

Figures label facilities by number.

| Facility | Data source | Photos | Mice | Sessions | Age range (months) | Strains | Females |
|---|---|---|---|---|---|---|---|
| 1 | INRCA (IT) | 928 | 692 | 9 | 3–34 | C57BL/6J, UM-HET3 | 69% |
| 2 | IRCAN (FR) | 711 | 210 | 5 | 1–28 | BALB/c, C57BL/6J | 78% |
| 3 | VivoArchitect (ES) | 624 | 157 | 4 | 2–27 | C57BL/6J | 17% |
| 4 | EPITERNA (ES) | 204 | 99 | 1 | 20–27 | C57BL/6J | 0% |
| 5 | IFOM (IT) | 195 | 65 | 1 | 7–12 | C57BL/6J, TERC<sup>−</sup>/<sup>−</sup> | 49% |
| 6 | ITP (US) | 100 | 50 | 1 | 5–21 | UM-HET3 | 50% |
| 7 | IOR (CH) | 84 | 19 | 2 | 2–10 | C57BL/6J | 0% |

### Image processing

Photographs were downscaled so that the longest side was at most 2,048 pixels. A YOLOv5n detector [17], fine-tuned from COCO-pretrained weights [26] on 247 manually labelled photographs (640-pixel input, batch size 8, up to 80 epochs with early stopping after 15 epochs without improvement), located the mouse, and its box, enlarged by 15% on each side, was cropped. Within this crop, Grounding DINO (base model, prompt “a mouse.”, box threshold 0.25) gave a bounding box that prompted SAM (ViT-H); of SAM’s three candidate masks, the one with the highest predicted overlap was kept. Detecting the mouse again inside the YOLO crop gave SAM a tighter box than the crop itself. The mask was reduced to its largest connected region with holes filled; pixels within a band of 2% of the mouse’s size along its edge were removed when their colour was within 12 CIELAB units of the local background or in the green-to-purple hue range of gloves and cloths. The background was set to flat grey (128), the mask edge softened inward by 1.5 pixels, and the mouse cropped with an 8% margin, padded to a square and resized to 512 pixels.

### Age model

Each crop was padded to a square with its corner colour, resized to 448 × 448 pixels and normalised with ImageNet statistics. A DINOv2 ViT-B/14 model pretrained by self-supervision on 142 million images (a vision transformer [27]; timm [28] model vit_base_patch14_dinov2.lvd142m) [20], with no further training, produced a 768-dimensional embedding; embeddings of the image and its horizontal mirror were averaged. A ridge regression on standardised embeddings (regularisation chosen by efficient leave-one-out cross-validation among 20 values from 0.1 to 10,000) predicted age. For the fine-tuning comparison, the network was trained on age with the same leave-one-facility-out splits, either end to end or starting from the ridge solution with the early transformer blocks frozen.

### Evaluation

The main split was leave-one-facility-out: each facility was predicted by a model fitted on the other six. A secondary split kept facilities in both training and test but held out mice (five-fold, grouped by mouse). Photographs of the same mouse in the same session were averaged before computing metrics. Within-session concordance was the fraction of pairs of mice in the same session with different ages for which the older mouse had the higher prediction (ties counted as half). 95% confidence intervals came from 200 bootstrap resamples of mice (1,000 for the comparison with human raters). Longitudinal concordance used all pairs of sessions of the same mouse. For calibration, predictions in a new facility were shifted by the mean error of k randomly chosen mice of known age, averaged over 200 draws, and evaluated on the remaining mice.

### Sex model and cropping comparison

The sex model used the same embeddings with a cross-validated logistic regression and the same leave-one-facility-out split; within-session concordance for sex was computed on male–female pairs (equivalent to a within-session area under the ROC curve). For the background comparison, the same age model was fitted on the YOLO crops with their background kept. Both comparisons were run at 224-pixel input, and their age results are compared only with each other.

### Photographic age gap

For each facility, a straight line was fitted to predicted against true age among control mice; the photographic age gap of a mouse was its predicted age minus the value of this line at its true age. This removes the regression towards the mean that makes young mice look older and old mice look younger, and the facility offset.

### Frailty, weight and survival

Frailty was assessed with a vitality score in one facility [21] (sign-reversed, so that higher means frailer) and a frailty index in another [4]. Within each facility, the score was regressed on the age gap and age (and sex where both were present), with standard errors clustered by mouse. In the frailty facility, mice that had received various treatments were included, and treatment was ignored. For pooling, scores were standardised within facility, signed so that higher means frailer, and combined by fixed-effect inverse-variance meta-analysis (heterogeneity I² [29]). Body weight was standardised within sex. Survival from the day of the photograph was analysed in the facility with the vitality score (the other facility with follow-up had 29 control mice) with Cox models [30] adjusted for age and sex, with robust standard errors clustered by mouse; sensitivity models added the vitality score or used only each mouse’s first photograph. Telomerase-deficient mice of each generation (G3, G4) were compared with the wild-type cohort photographed alongside them by regression of the age gap on group and age, with robust standard errors.

### Sex and strain

Within-session concordance was computed separately within each sex and strain (pairs of the same group only). Differences in age gap between groups were computed within each facility containing both groups, as differences of per-mouse mean gaps with bootstrap confidence intervals; facility estimates were pooled weighting by number of mice.

### Human raters

100 photographs from facilities not used for training, stratified across five age bands and including pairs from the same session, plus 10 hidden repeats, were shown in random order through a web form. Raters entered an age in months and received no feedback. Scores were computed exactly as for the model on the same photographs, with paired bootstrap differences.

### Occlusion maps and body regions

For each photograph, the age model fitted without its facility was used. A square window of one-eighth of the image side was moved in steps of half its width and replaced by flat grey; the change in predicted age was averaged over all windows covering a pixel. Body regions were defined from the mouse mask: thin parts removed by morphological opening were assigned to the tail (the longest, most elongated part) or limbs; the main body axis was found by principal components; the end joined to the tail was the rear. Head and rear were the outer quarters of the body length, mid-body the rest. Mice without a visible tail were left unoriented and excluded from head and rear comparisons. Region effects were summarised as mean change per pixel, with confidence intervals from resampling facilities.

### Data availability

The data supporting this study are available on Zenodo (https://doi.org/10.5281/zenodo.23689447) under a CC BY 4.0 licence: the 2,846 photographs as background-removed and detector crops, with facility, session, mouse identifier, age, sex, strain and training status for each; the frailty, vitality, body-weight and survival data; the human-rater task and anonymised answers; the photographs and labels used to train the mouse detector; and the trained detector and age models. Facilities are identified by number, as in the figures (key in the Methods table).

### Software and code availability

Python 3 with PyTorch [31], timm [28], transformers [32], scikit-learn [33], statsmodels [34] and lifelines [35]. Code, trained models and figure scripts are available at https://github.com/perez-kevin/mouse-photographic-age.

## Acknowledgements

We thank Andrea Amoroso (Charles River Laboratories, Calco, Italy) for precious support in mouse management. We thank Alberto Parras, Florinda Melendez (VivoArchitect, Switzerland), Richard Miller (ITP) and team members (EPITERNA) for providing photographs of mice.

## Author contributions

K.P. conceived and designed the study, performed the analyses and wrote the manuscript. F.R. and A.M. contributed to the analysis. C.M., L.T., M.S., G.B. and S.M. contributed to the experimental work, animal care and data collection. L.T., K.Pa. and M.S. estimated mouse ages from photographs as human raters. M.M. provided the frailty and survival data. F.d’A.d.F., A.A., D.B. and M.M. provided mice and photographs from their facilities and supervised the work there. K.Pa. reviewed the manuscript and advised on collaborators. All authors read and approved the final manuscript.

## Ethics statement

At INRCA (Facility 1), photographs were obtained exclusively from untreated or control mice included in research projects conducted at the INRCA Geriatric Mouse Clinic, approved by the Italian Ministry of Health (authorizations No. 1074/2020-PR, 87/2022-PR and 392/2019-PR) or conducted under the applicable procedure (scientific project No. 8502E.N.4VV). At VivoArchitect (Facility 3), mice experiments were conducted under protocol no. 15012/2024/024 approved by Xunta de Galicia. At EPITERNA (Facility 4), mice experiments were conducted under protocol no. 2023/160204 approved by Xunta de Galicia. At IFOM (Facility 5), experiments were authorized by the Italian Ministry of Health, the national Competent Authority (project number 565/2021). For the ITP (Facility 6), the project was approved by the University of Michigan Institutional Animal Care and Use Committee (IACUC). At the other facilities, all procedures involving animal observation and experimentation were reviewed and approved by the appropriate institutional ethics committee.

---

**Supplementary Fig. 1.**
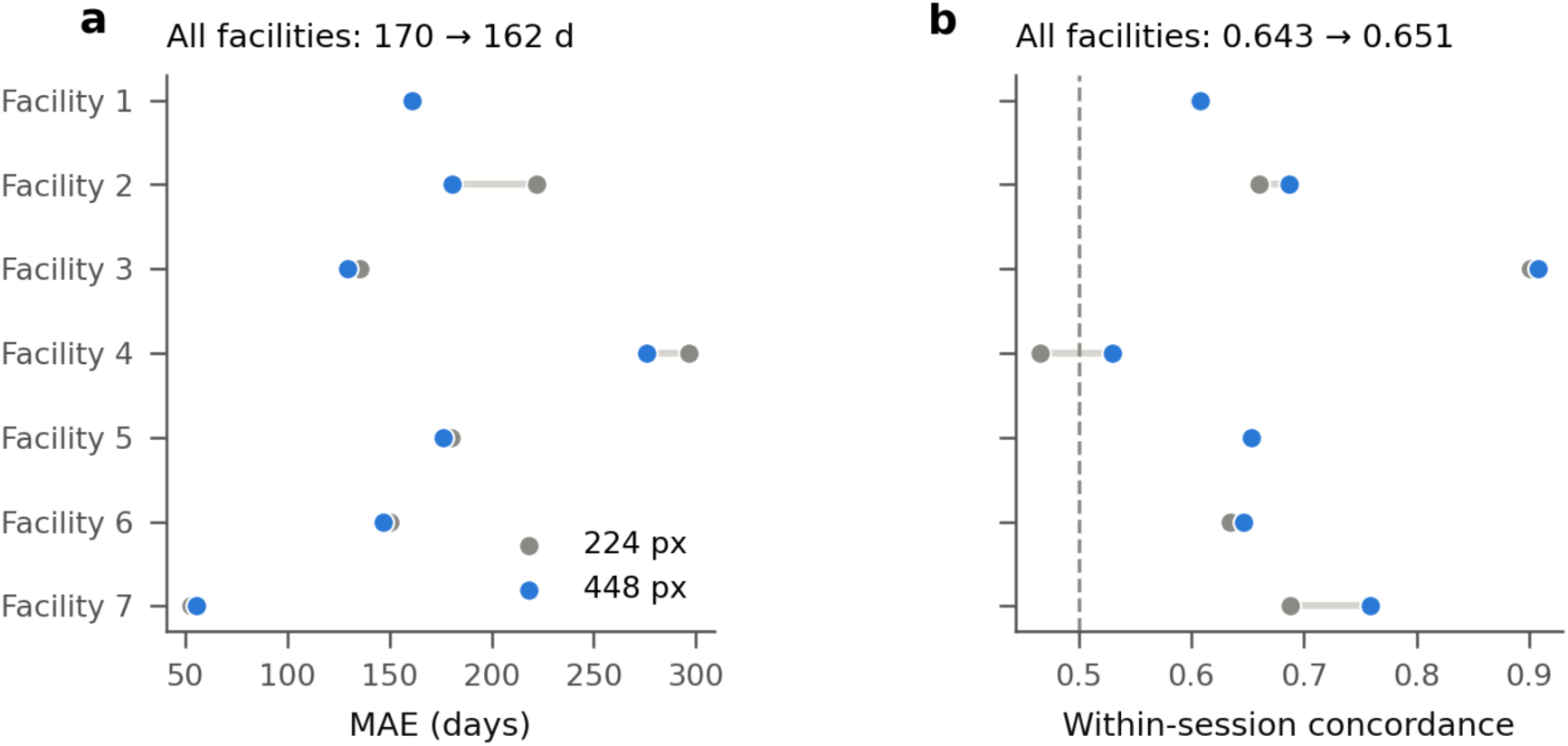
Input resolution. Error (a) and within-session concordance (b) per facility for 224- and 448-pixel input.

